# PLO3S : Protein LOcal Surficial Similarity Screening

**DOI:** 10.1101/2022.03.25.484718

**Authors:** Lea Sirugue, Florent Langenfeld, Nathalie Lagarde, Matthieu Montes

## Abstract

The study of protein molecular surfaces enable to better understand and predict protein interactions. Different methods have been developed in computer vision to compare surfaces that can be applied to protein molecular surfaces. The present work proposes a method using the the Wave Kernel Signature : Protein LOcal Surficial Similarity Screening (PLO3S). The descriptor of the PLO3S method is a local surface shape descriptor projected on a unit sphere mapped onto a 2D plane and called Surface Wave Interpolated Maps (SWIM). PLO3S allows to rapidly compare protein surface shapes through local comparisons to filter large protein surfaces datasets in protein structures virtual screening protocols.

## 1 Introduction

Proteins are central in most biological processes. Proteins can be described through their sequence, structure, surface and/or function(s). The protein surface is an abstract, geometric representation of the protein potential interactions, structure, fold, and sequence [11, 15, 30]. Proteins sharing a related function display similar surfaces that can be independent of their sequence and/or structure similarity [15, 23, 39]. Different methods based on protein surface comparison have been developed for protein-protein interactions prediction [11, 34, 40], protein structural alignment [25] or protein shapes classification [8, 12, 13, 15, 23, 24, 27, 38].

Surface comparison methods can be classified into different categories depending on their surface shape descriptor computed from the surface. (1) The methods based on spectral geometry establish a relationship between the surface shape and the spectra of the Laplace-Beltrami operator. A spectrum of the Laplace-Beltrami operator is a fingerprint composed of the eigenvalues obtained using the differential Laplace-Beltrami operator [2, 32, 44]. (2) The methods based on histograms summarize local or global geometrical or topological features of the surface [17, 35, 37, 45]. (3) Projection-based methods use the projection(s) of the protein topography in the 2D space [7, 29]. (4) Zernike-based methods use the moments of 3D Zernike polynomials [5, 19]. They have been widely used on protein surfaces and display high performances in retrieval [5, 14, 18, 19]. (5) The last category comprises methods based on geometric learning using convolutional neural networks [11, 26].

Surface shapes can be described globally or locally. A global surface shape descriptor describes the surface shape of the whole object [17, 31, 36] which allows direct comparisons of the whole surface shape of different objects whereas a local surface shape descriptor is defined over a surface patch and allows comparisons with other surface patches.

To our knowledge, no spectra-based method has yet been developed for protein surface comparison. In the present work, we describe Protein LOcal Surficial Similarity Screening (PLO3S), a fast protein surface shapes comparison method based on a new local spectral descriptor, Surface Wave Interpolated Maps (SWIM). SWIM is a wave kernel signature (WKS [2]) conformally projected on a 2D plane. In PLO3S, the values of SWIM are processed using a dense point-to-point comparison. PLO3S is designed to blindly screen large protein surfaces datasets in order to discard protein surfaces that do not share high local surficial similarity to the query. This allows for (1) further protein surface shapes screening with finer local surface shape comparison methods that cannot handle large datasets and (2) reducing the number of false positive potential binding sites in the context of target fishing for adverse interaction screening or poly-pharmacology in a drug discovery pipeline or protein-protein interactions annotation.

## 2 Materials and methods

### 2.1 PLO3S

Protein LOcal Surficial Similarity Screening (PLO3S) is a screening method for finding local and surficial similarity of proteins, independently of the sequence of the proteins. PLO3S is described in two steps: (1) the computation of the Surface Wave Interpolated Maps (SWIM) descriptor and (2) the computation of the surface shape similarity

#### Computation of the SWIM descriptor

The Solvent Excluded Surface of the input protein structure (Figure 1a) is computed using EDTSurf [47] (Figure 1b). The Wave Kernel Signature (WKS) is then computed on the resulting 3D mesh *M* [2] (Figure 1c). For each point of the surface of the 3D mesh, a vector of size *N*, representing the WKS descriptor is computed. This descriptor, based on the eigenvalues of the Laplace-Beltrami operator, has the property of invariance to isometry and is robust to perturbations [2].

**Figure 1:**
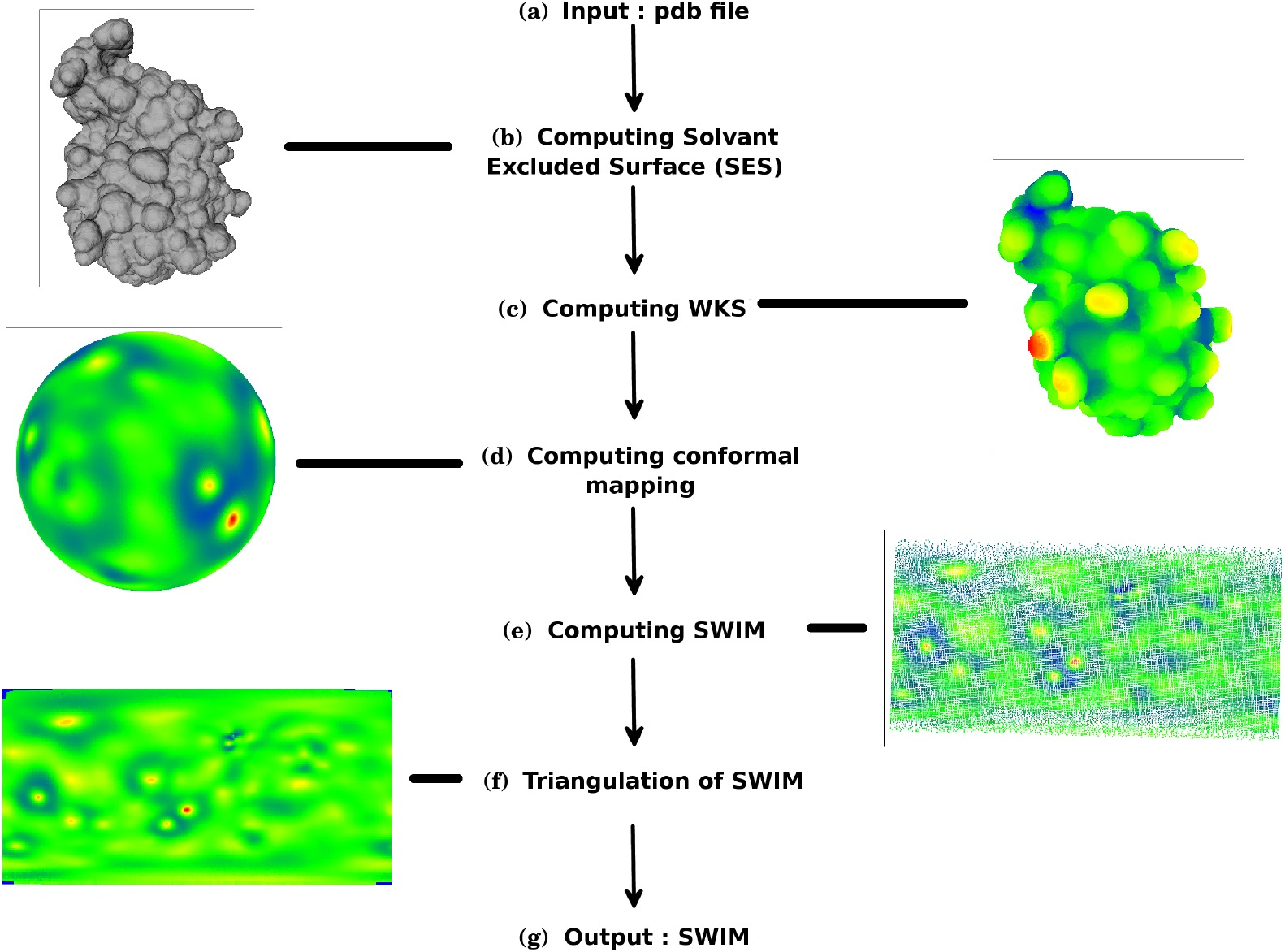
Diagram for the generation of a SWIM. The SES is computed using EDTSurf [47] (a)(b). In a second step, the Wave Kernel Signature (WKS) [2] is calculated for each point of the surface (c). The surface is then projected on a unit sphere [1] (d). The sphere is mapped onto the 2D plane [8] (e). The points of the map are interpolated (f) to form the final descriptor called SWIM (g).

On a second step, the 3D mesh is projected on a unit sphere *S* as described by Angenent et al. [1] (Figure 1d). The frame of the unit sphere is defined using a reference point on its surface called the *pole*. This transformation is conformal and bijective. It is to note that even if the distances and surface areas are not preserved in the projection, they are only modified by a scaling factor.

Then, the unit sphere is transformed onto the 2D plane based using the two spherical coordinates of the angles (*θ*, *ϕ*) [7, 8] (Figure 1e). A map of size (*θ*_*max*_ − *θ*_*min*_)/*δ,* (*ϕ*_*max*_ − *ϕ*_*min*_)/*δ* is created. *θ*_*max*_ and *θ*_*min*_ are the maximum and minimum values of *θ* and *ϕ*_*max*_ and *ϕ*_*min*_ are the maximum and minimum values of *ϕ*. *δ* is the step for dividing the sphere in areas, where each area are represented by a point on the map in the discrete plan. The value associated to each point on the map is the WKS descriptor represented by a vector (Figure 2). These maps are called Surface Wave Interpolated Maps (SWIM).

**Figure 2:**
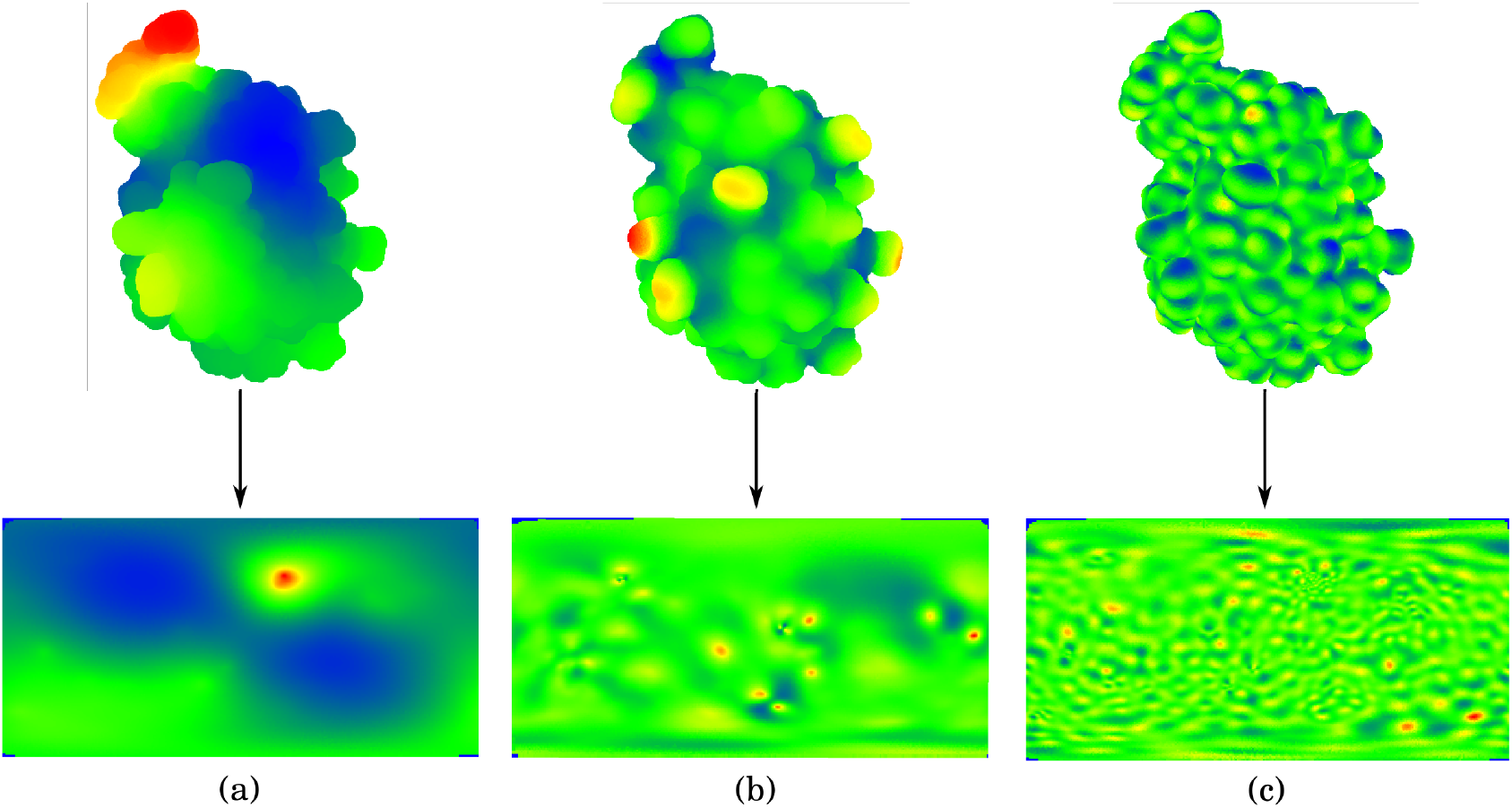
WKS on the surface of one conformation of ubiquitin (5xbo A 1) (top) and its corresponding SWIM (bottom) for the 1st (a), 50th (b) and 100th (c) value of the WKS.

The final step is the interpolation of the point of the map (Figure 1 f). The map is encoded as an image and the projection on the 2D plane is not filling e ach p ixel w ith a v alue. I t c an c reate a n unbalance if a map has large areas with no value, or if the neighborhood of a point of the map is considered for comparison. So, for each pixel with no value, the three points on the map defining the triangle with the smallest area containing the pixel are used for interpolating the value of this pixel.

The main issue with this representation is the deformation in the neighborhood of the poles while passing from the unit sphere to the 2D plane. To handle this issue, we use a multi-view approach where the pole axis is rotated by an angle *α* in the planes perpendicular to each of the three Cartesian axis of the unit sphere (Figure 3). Then, a SWIM is created as above-mentioned (Figure 1g). We uses 7 projections with a 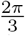 rotation (Figure 3) to generate a set of SWIMs as it represents the best trade-off between the deformation at the poles and the computational efficiency.

**Figure 3:**
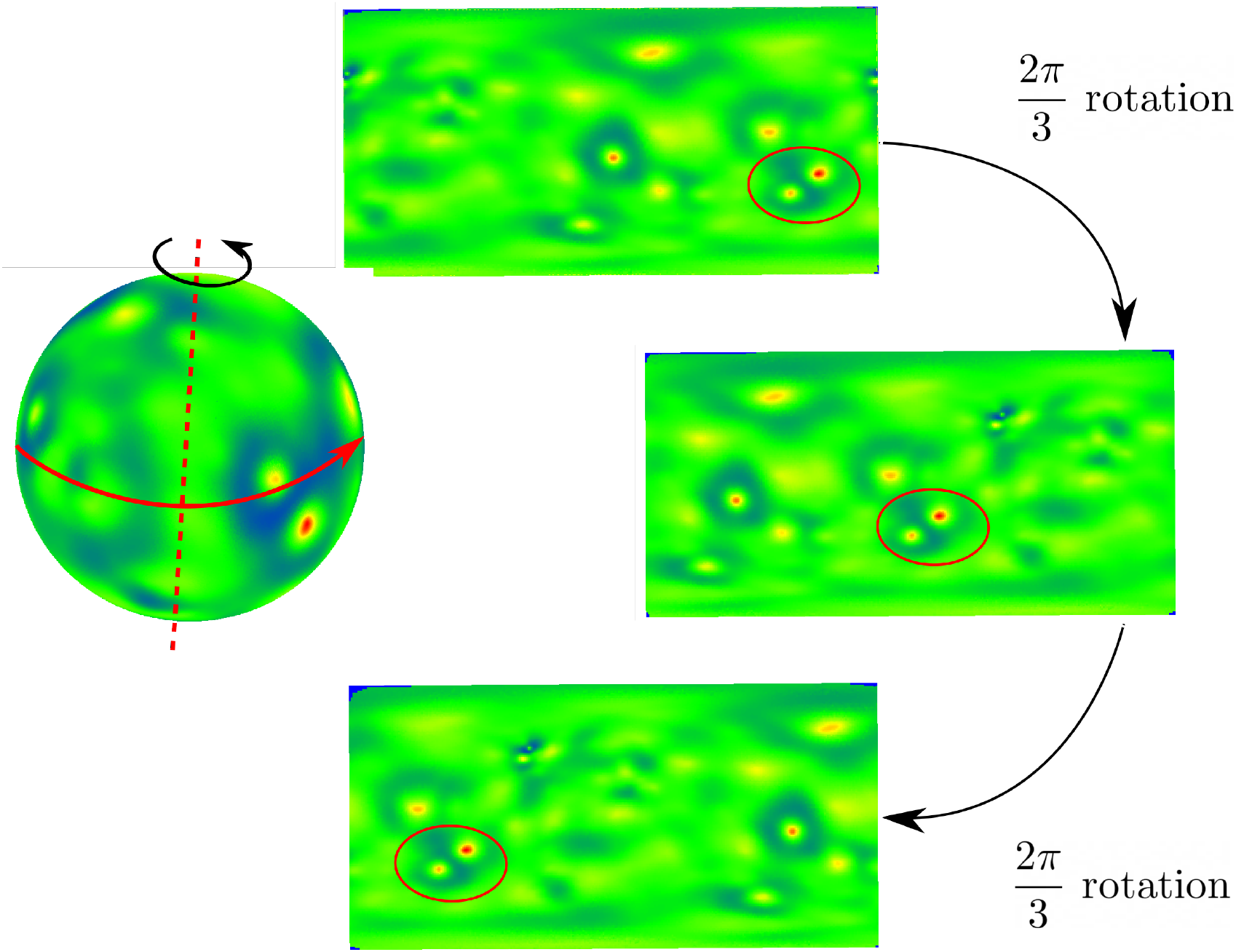
Unit sphere (Left Panel) and its corresponding SWIMs for the 50th value of the WKS with a rotation of 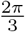 around a pole of the unit sphere of Ubiquitin (5xbo_A_1).

#### Computation of the surface shape similarity

A dense exhaustive point by point comparison is performed between the generated SWIMs.

Two shapes *T* and *V* are compared by searching the best matches of the vectors of their respective SWIM *C*_*T*_ and *C*_*V*_. To compare the two vectors *H*_*kT*_ and *H*_*kV*_ at points *k*_*T*_ and *k*_*V*_ from the maps *C*_*T*_ and *C*_*V*_, respectively, the Earth Mover’s Distance (EMD) [33] is used. EMD is a suitable distance for WKS because it takes into account the proximity of the bins of an histogram and the sequence of the eigenvalues of the WKS represents in decreasing order a surface scale of the region on the surface which can be assimilated to an histogram.

Since the WKS is represented as a 1D array, the EMD equation can be simplified as the sum of the absolute differences between the cumulative values of the WKS.

Given two WKS, two vectors with weights equal to one, *X* = {*x*_1_, …*x*_*n*_} and *Y* = {*y*_1_, …*y*_*n*_}, ∀ *i, j* ∈ [1..*n*] *d*_*ij*_ = |*x*_*i*_ − *y*_*j*_ the *L*_1_ distance is defined and a function *f*_*ij*_ such as it minimizes the following equation :

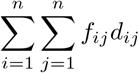

with the constraints *f*_*ij*_ > 0, 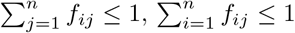 and 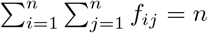.

The WKS is represented as a 1D vector with no weights, which means that the EMD equation can be simplified as the sum of the absolute difference between the cumulative values of the WKS. Given 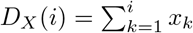 and 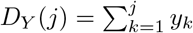, then the EMD *L* between two WKS *X* and *Y* is defined as :

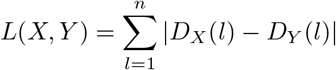

The surface shape dissimilarity score for the shapes *T* and *V* is the sum of the best distance of each point *L*. If *C*_*T*_ is composed of *N*_*T*_ points and *C*_*V*_ of *N*_*V*_ points, then the score *S*(*T, V*) of dissimilarity between *T* and *V* is :

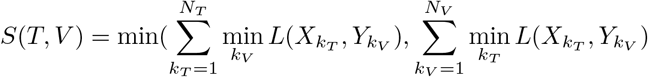

The surface shape dissimilarity score is normalized between 0 (identity) and 255 (maximum dissimilarity of the shapes).

**GPGPU optimization** The computation of the shape similarity is performed in parallel using the General Purpose GPU (GPGPU) sum reduction technique [6, 9]where the data is divided into fixed sizes that fit the internal memory of the GPU.

### 2.2 Spectral geometry based shape comparison methods

**Heat Kernel Signature** [44] is derived from the Heat Kernel which represents the diffusion of heat on an object as a function of time. The HKS is based on the spectrum of the Laplace-Beltrami operator. HKS is a local descriptor with the property of isometric invariance and is stable against perturbations [44].

**Wave Kernel Signature** [2] is based on the spectrum of the Laplace-Beltrami operator. It describes the energy of quantum particles on the surface of the shape based on the wave equation which is a solution of the Schrödinger’s equation. A signature is created with the solution of the wave equation. WKS is a local descriptor with the property of isometry and scale invariance. Contrary to HKS, in WKS, time is not taken into consideration. It is replaced by the energy of the particle. For the WKS, the energy is related to the size of the geometry; a large energy represents a local geometric feature while a small energy represents a global geometric feature.

### 2.3 Protein structure comparison methods

In **Combinatorial Extension (CE)** [43], the protein is represented as a set of fragments of 8 amino-acids. A fragment is aligned to another fragment composed of at least 8 amino acids and form an Aligned Fragment Pair (AFP). Using constraints on the maximum distance between AFPs, the AFPs are assembled to form a longer path. Then, an optimization is performed using the Z-score and dynamic programming [28].

In **TM-Align** [49], the protein structure is aligned independently of sequence length using TM-score [48]. In a first step, an alignment based on dynamic programming is proposed. It is decomposed into a residue-to-residue alignment and a secondary structure alignment. In a second step, the TM-score evaluates the matrix score of the alignment. The proteins are aligned again to increase the score, and this step is repeated until stability is found.

In **DeepAlign** [46], the proteins are aligned according to spatial proximity, evolutionary links and hydrogen bond similarity. The score is a combination of a value for amino-acids substitution [16], conformation substitution formed by the angles of the pseudo-bonds of the *C*_*α*_ atoms [50], the TM-score to estimate spatial proximity, and hydrogen bond similarity.

### 2.4 Benchmarking Dataset

The benchmarking dataset is a subset of the Protein Shape Retrieval Contest track dataset of SHREC19 [20]. The subset contains 14 protein shape classes, based on the protein level of the SCOPe classification [10]. Each protein shape class contains between 20 and 30 conformations for a total of 403 protein conformations. The IDs used are in the following format : PDBID_A_X, “PDBID” being the id of the PDB with 4 letters, “A” being the protein chain and the “X” being the conformation number according to the PDB file. For each structure of the dataset, the Solvent Excluded Surface (SES) is computed using EDTSurf [47] with default parameters. EDTSurf output triangular meshes are stored as .ply file, converted to .off and .pcd formats, required by the different shape comparison methods.

### 2.5 Performance evaluation metrics

The performance in retrieval of each method are evaluated using Precision-Recall and negative predicted value curves. The Precision-Recall plot draws the recall *R* as a function of the precision *P*. Precision *P* is the ratio of targets from class *C* retrieved within all objects attributed to class *C*, while recall *R* represents the ratio of retrieved targets from class *C* compared to |*C*|, the size of class *C*. Precision-recall curves are computed using the Princeton Shape Benchmark tools [42]. The negative predictive value evaluates the percentage of objects rightfully classified as negative within the negatives.

### 2.6 Runtime

All calculations were performed on a 64-bit Linux Ubuntu desktop computer with an Intel Xeon 2.30GHz CPU, 32GB of RAM and a Quadro K4200 4GB GPU.

## 3 Results

The evaluation of the performance of PLO3S in enrichment has been performed on a protein shapes dataset derived from the Protein Shape Retrieval Contest of the SHREC 2019 community benchmark [20]. The performance of PLO3S is compared with spectral, geometry-based shape comparison methods and protein structure comparison methods. For all the shape comparison methods, blind all-to-all dense comparison have been performed on the whole dataset. Dense comparison is performed when all-to-all descriptor points are compared during the computation of the surface shape similarity.

### 3.1 Illustration of PLO3S method with selected examples

Three proteins were selected from the SHREC19 dataset, herein used as the benchmarking dataset, to illustrate the PLO3S method functioning and outputs. The selection was made to include two proteins presenting similar SWIMs, ubiquitin and thioredoxin, and one protein presenting a dissimilar SWIM compared to the two former proteins, Macrophage Inflammatory Protein (MIP, Figure 4). Ubiquitin and thioredoxin display a globally spherical shape whereas MIP is more elongated. The SWIMs of MIP are less dense, as illustrated by the large blue area representing values of the WKS close to 0. The area containing more discriminating information, the red area, is located at the bottom of the SWIM (close to the pole) and then distorted because of the mapping of the unit sphere to the 2D plane.

**Figure 4:**
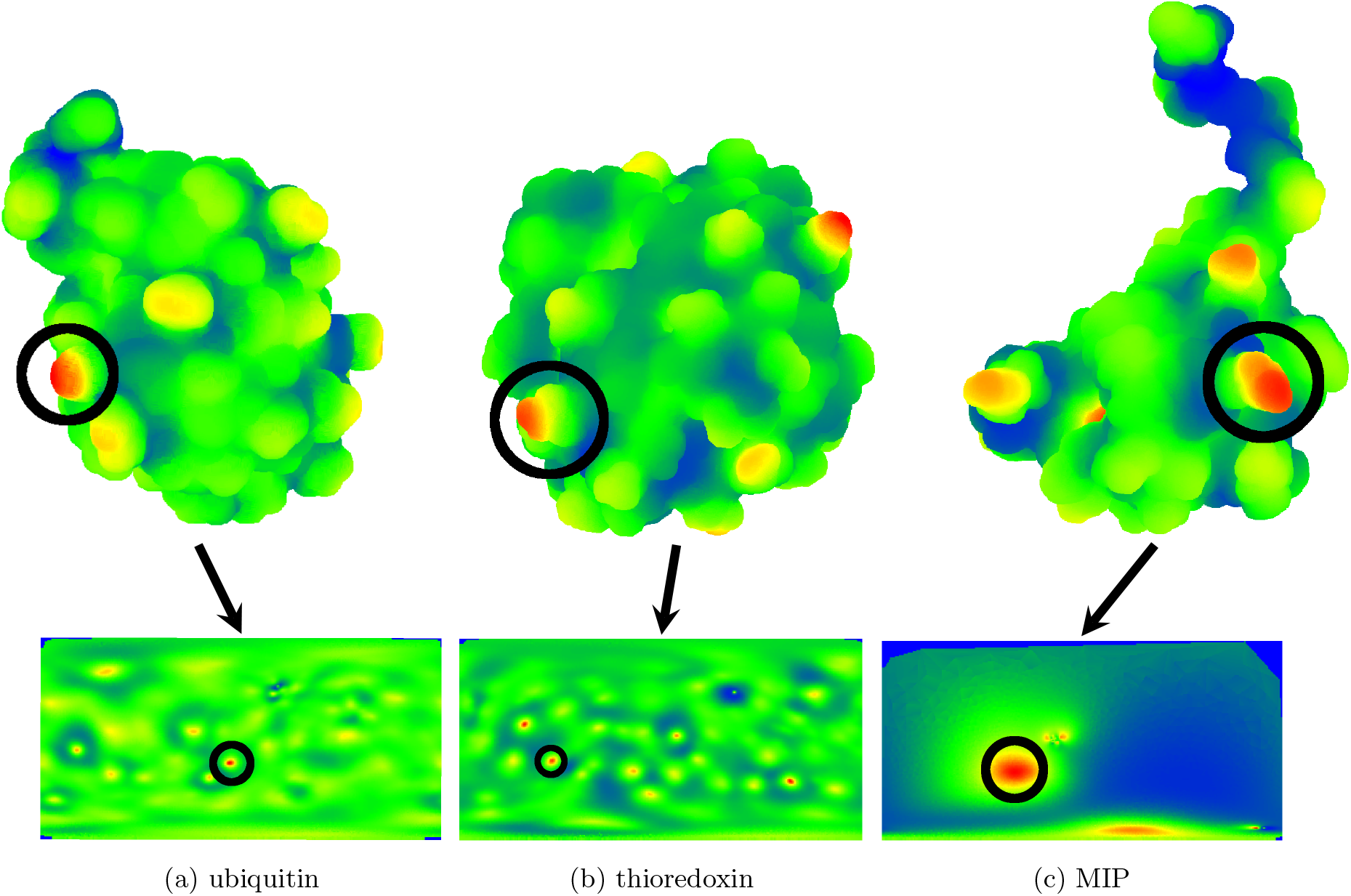
Illustration of the 50th value of the WKS on the SES’ (Upper Panel) and their corresponding SWIMs (Lower Panel) for (a) ubiquitin 5xbo_A_1, (b) thioredoxin 1trv_A_1 and (c) Macrophage Inflam-matory Protein 1hun A_1.

The dissimilarity scores between different conformations of ubiquitin, thioredoxin and MIP are presented in Figure 5. The ubiquitin and thioredoxin are very similar, as evidenced by dissimilarity scores ranging from 0 to 40. On the contrary, the dissimilarity scores of MIP, compared to ubiquitin or thioredoxin, varies from 30 to 255.

**Figure 5:**
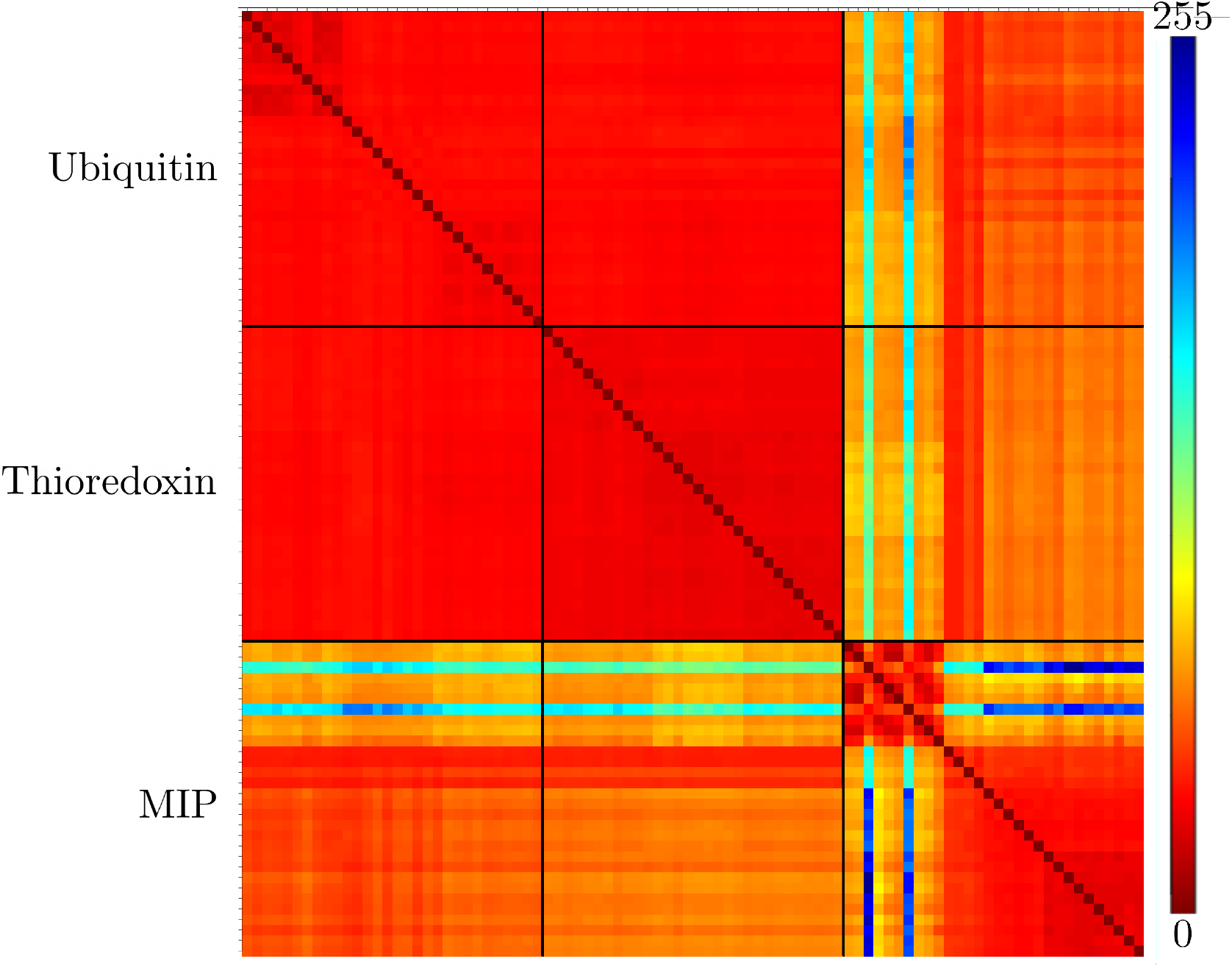
Matrix of dissimilarity scores of PLO3S method on 3 selected examples: ubiquitin, thioredoxin and Macrophage Inflammatory Protein (MIP)

Within the MIP class, the dissimilarity scores are separated into two groups of respectively 10 and 20 conformations. Dissimilarity scores within these two groups range from 0 to 50 and are similar to intraclass scores of ubiquitin and thioredoxin (see the lower right corner of Figure 5, which exhibits two red squares corresponding to the two MIP groups). The score between these two groups are significantly different, ranging from 60 to 255. The score of the third and seventh conformations (1hun A 3, 1hun A 7) are even more distinct with scores varying from 120 to 255.

The first ten MIP conformations have a more cylinder-like shape (Figure 6a and 6b) than the last twenty conformations (Figure 6c).

**Figure 6:**
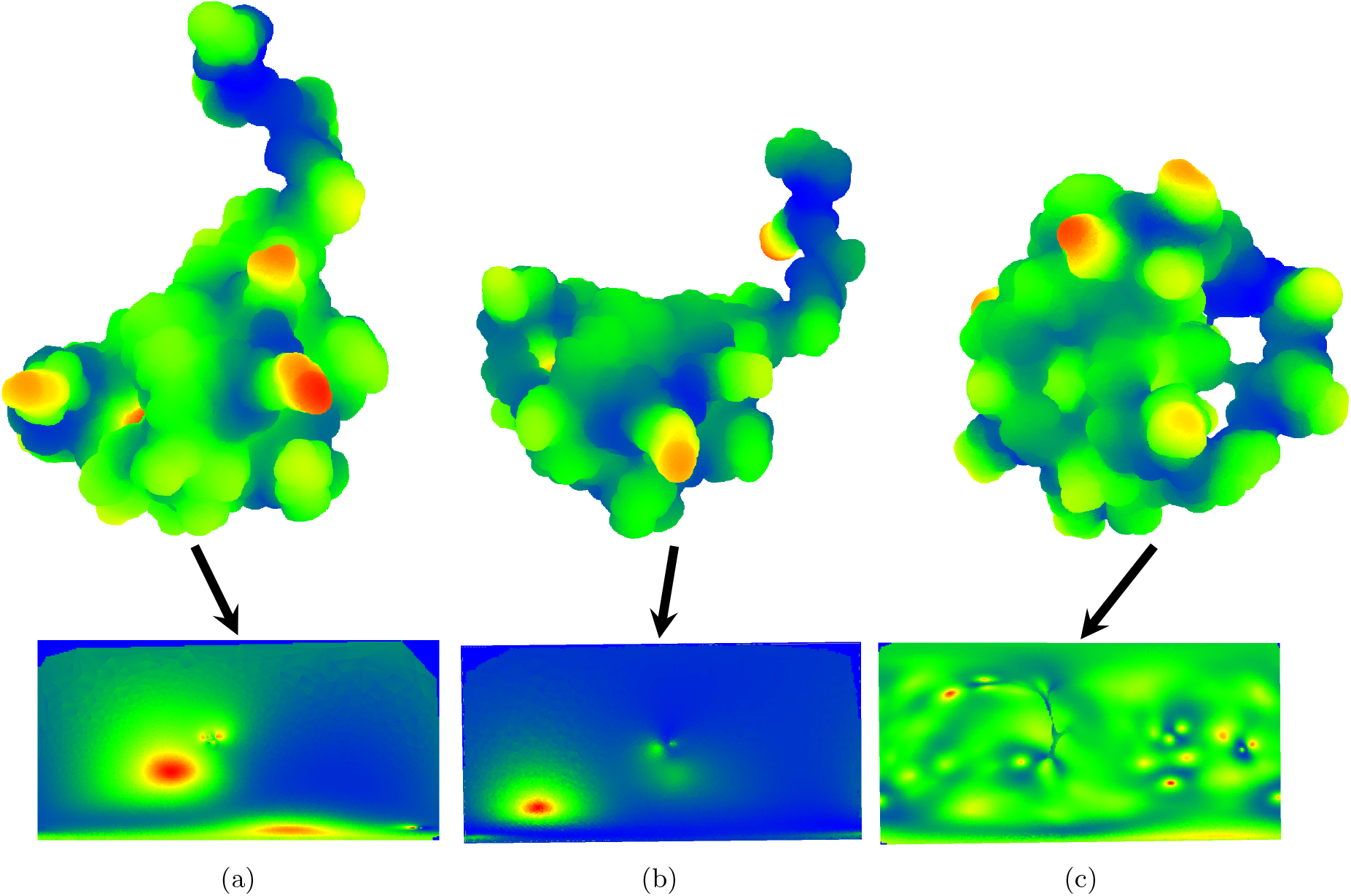
Illustration of the 50th value of the WKS on the SES’ (Upper Panel) and their corresponding SWIMs (Lower Panel) for three structures of the Macrophage Inflammatory Protein (MIP) : (a) 1hun_A_1, (b) 1hun_A_3 and (c) 1b50_B_1.

Similar to intra-class scores, the first ten MIP conformations have a significantly high dissimilarity score with all other conformation, varying from 60 to 255 and the score of the third and seventh MIP conformations (1hun_A_3, 1hun_A_7) ranging from 120 to 255.

### 3.2 Evaluation of PLO3S in enrichment

The overall results for PLO3S are shown in figure 7. Precision decreases steadily from 0.9 to 0.2 for the PLO3S method (Figure 7a) as the recall increases from 0.05 to 1. The recall increases up to 0.9 when considering the first 62% of the results, and then increases steadily up to 1. This indicates a compromise between downsizing the dataset and removing a maximum of true negatives. Negative Predictive Value (NPV) ranges from 0.93 to 0.99 with a peak at 37% of the dataset size (Figure 7c). This peak corroborates the recall value stabilizing at 62% of the dataset size as the NPV is determined by the negatives while the recall is based on the positives.

**Figure 7:**
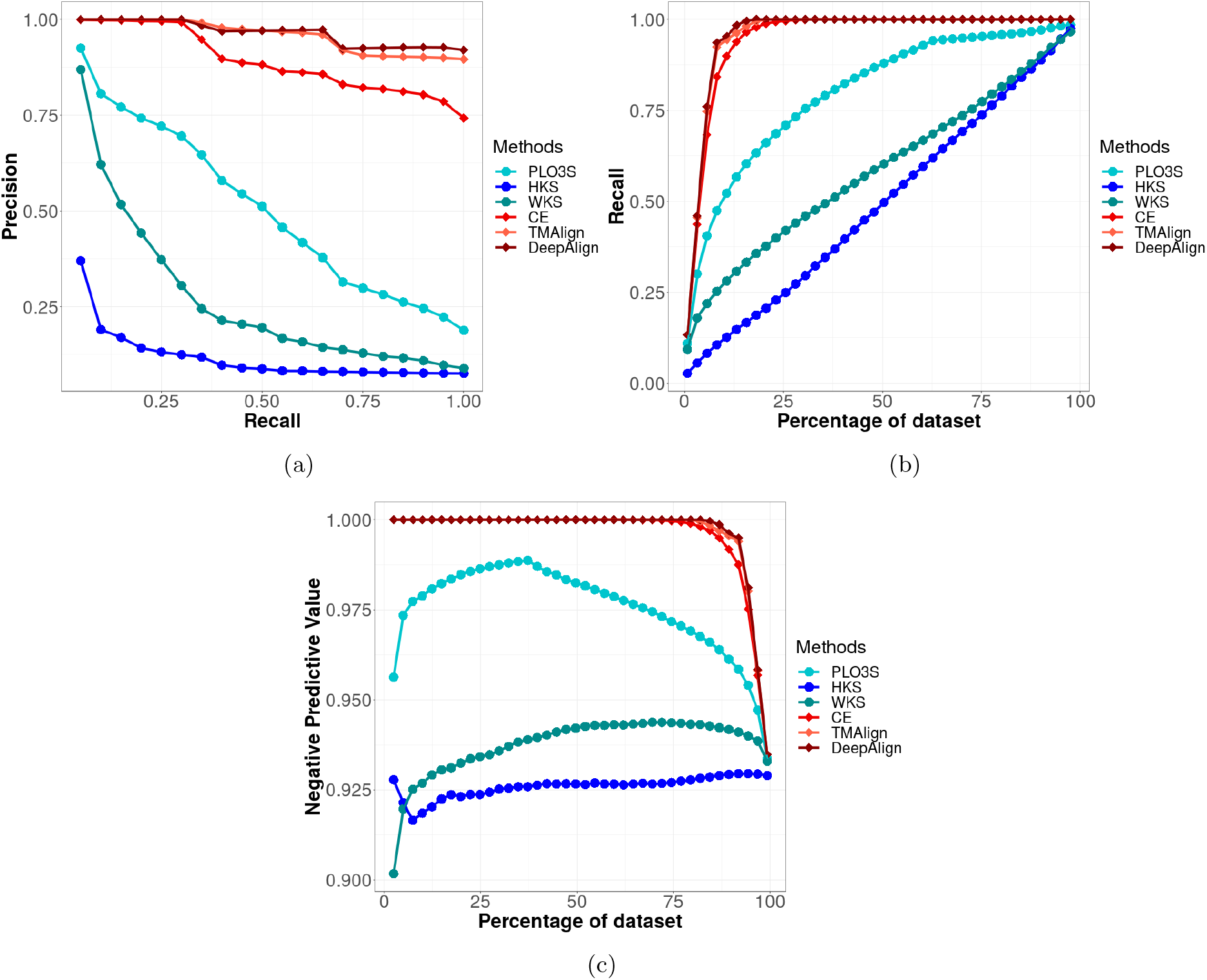
a Precision-Recall curves for PLO3S, HKS, WKS, CE, TM-Align and DeepAlign. b Recall values for different t hresholds f or p ositives. c N egative P redictive V alues f or d ifferent th resholds for negatives.

The computation time of the descriptor SWIM and of the comparison of two SWIM descriptors are shown in Table 1. On average, 23 minutes and 35 seconds are necessary to generate a SWIM. The average computational time associated with the comparison of two SWIM descriptors is 0.7 seconds.

**Table 1:**
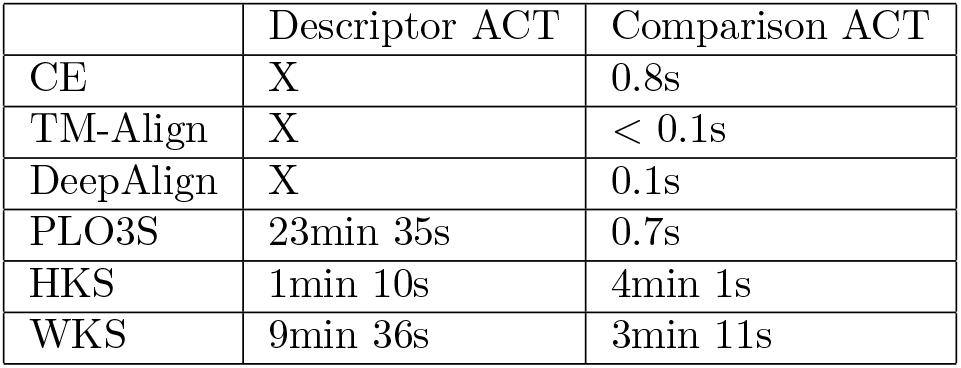
Average computation time (ACT) for one descriptor and comparison for CE, TM-Align, DeepAlgin, PLO3S, HKS and WKS in seconds.

### 3.3 Comparative evaluation of the performance of PLO3S in enrichment with spectral geometry based shape comparison methods

We compared our PLO3S method with 2 other spectral geometry based methods for shape comparison, WKS and HKS which have already shown good performance [21, 21, 22].

The PLO3S method is the top performer for the precision-recall metric (Figure 7a). The precision for WKS and PLO3S is about 0.9 for a 0.05 recall, while the precision of HKS is 0.37. The precision of WKS and HKS methods is lower than 0.05 for a recall of 1 while the PLO3S precision is superior, around 0.2. Both HKS and WKS descriptors are computed and compared on 3D surface. However, PLO3S descriptor, called Surface Wave Interpolated Maps (SWIM), is based on a 2D space. Despite having one less dimensional space, SWIM shows superior performance to the HKS and WKS descriptors.

The PLO3S method has the highest recall compared to HKS and WKS for all sizes of the positives set (Figure 7b). The recall curve of PLO3S increases rapidly at the beginning and stabilizes around 62% of the dataset size, while the recall of HKS and WKS increases steadily.

The PLO3S method outperforms both HKS and WKS methods for Negative Predictive Value (NPV) (Figure 7c). NPV ranges from 1 to 0.9, as the size of the true negatives set is approximately 13 times larger than the size of the true positives set. The NPV for PLO3S ranges from 0.93 to 0.99 and its maximum is at 37% of the dataset size. On average, HKS and WKS show stable NPV, around 0.93 for WKS and 0.92 for HKS which is globally inferior to PLO3S

We determined the average descriptor computation time and comparison time in Table 1 for the three methods. The descriptor computation time includes the processing time from the input mesh to the final descriptor. PLO3S has the slowest descriptor computation time with 23 minutes and 35 seconds, while the fastest descriptor computation time is 1 minute and 34 seconds (HKS). On the contrary, the fastest comparison time is 0.7 seconds for PLO3S, while the slowest is HKS with 4 minutes and 1 second on average.

### 3.4 Comparative evaluation of the performance of PLO3S in enrichment with protein structure comparison methods

The 3 standard protein structure comparison methods, CE, TM-Align and DeepAlign outperformed the PLO3S method in all three measures, precision-recall, recall and negative predictive value (Figure 7). While the precision of PLO3S is about 0.9 for low recall and 0.2 for high recall, DeepAlign and TM-Align have higher overall precision, ranging from 0.89 to 1 for different associated recall.

The PLO3S recall stabilizes around 0.9 for a dataset size of 62%. The recall of protein structure comparison methods is 1 after the first 20% of the results.

The PLO3S NPV is inferior to the protein structure comparison methods, with its highest NPV of 0.99 at 37% of the dataset size. The NPV of the protein structure comparison methods equals 1 for a 2% to 75% dataset size and reaches 0.93 for 100% of the dataset.

The protein structure comparison methods do not require the computation of a descriptor because the comparison and scoring parts are directly based on the structure (Figure 1). The comparison time is 0.8 second, 0.7 second, 0.1 second and inferior to 0.1 second for CE, PLO3S, DeepAlign and TM-Align respectively. The comparison time of PLO3S is thus similar to the comparison times of the protein structure comparison methods.

## 4 Discussion

### 4.1 Illustration of PLO3S method with selected examples

In order to illustrate the PLO3S method functioning and outputs, three proteins, ubiquitin, thioredoxin and MIP, were selected and further analyzed. The SWIMs of ubiquitin and thioredoxin are similar, which is corroborated by low values of their corresponding dissimilarity scores. Conversely, the high values in dissimilarity scores obtained by comparing the SWIMs of MIP and ubiquitin and thioredoxin illustrate their difference in shape since MIPs display an elongated shape.

An elongated shape, not easily projectable into a unit sphere, may be prone to more noise brought by the projection.

This can be observed with the differences between the scores of the group of 10 conformation of MIP opposed to the other 20 conformations of MIP. The first group of 10 MIP conformations has more elongated shapes (Figure 6a and Figure 6b) compared to the second group of 20 MIP conformations (Figure 6c).

Interestingly, for the MIP, the dissimilarity scores highlighted the existence of 2 distinct groups of conformations. All these conformations were gathered under the same MIP protein label by the SCOPe classification, but when investigating further we noticed that these 2 groups of conformations belong to similar yet distinct proteins. The 10 first MIP conformations are from the Macrophage Inflammatory Protein 1 alpha (MIP-1alpha), and the 20 last MIP conformations are from the Macrophage Inflammatory Protein 1 beta (MIP-1beta). The conformations available in the data set for these 2 proteins present different shape and thus different surface topology. Even if MIP-1alpha and MIP-1beta display 68% sequence identity, PLO3S allowed to highlight their surficial dissimilarity.

All the dissimilarity scores of the first group of 10 MIP conformations are high compared to all other scores, indicating the noise sensitivity of an elongated shape projected onto a unit sphere. In particular, the third and seventh conformations of MIP (1hun_A_3 and 1hun_A_7) show significantly higher scores than the other conformations of the 10 MIP conformations.

Because the scores for the third and seventh MIP conformations are higher in all respects than the other, they indicate an outlier probably caused by the elongated shape not fitting well on the unit sphere.

### 4.2 Evaluation of PLO3S in enrichment

In order to evaluate the PLO3S performance in shape retrieval, we used a benchmarking data set with 403 protein conformations for 14 protein shape classes. The performance of the PLOS3S method was measured using Precision-Recall and negative predicted value curves. Around a threshold of 62% of the dataset size, the recall of the PLO3S method reaches a value of 92% The highest value for NPV is around 38% which means that further analysis of the last 38% of the dataset is not relevant. Thus, the dataset size can be reduced by 38% (while discarding less than 10% of the real positive objects). This allows to dramatically reduce the effort required to screen a large dataset in a hierarchical protocol. The performance of PLO3S in terms of recall and NPV indicates that our method meets its main objective: selecting most of the positive objects while discarding a large number of negative objects with high confidence to safely decrease the size of very large datasets in a context of protein surface shapes screening.

The average computational time required to compute the SWIMs, the PLO3S descriptors, accounts for most of the time of the method. This preprocessing step is performed once for a given object and the resulting SWIMs can be stored for later comparisons/screens in a SWIM database describing protein surface shapes. Here, the critical aspect for a large database screeening is the computational time required to compute the shape dissimilarity between two protein shapes which is satisfactory (0.7 seconds in average).

### 4.3 Comparative evaluation of the performance of PLO3S in enrichment

Spectral geometry through the spectra of the Laplace-Beltrami operator is a common approach in the field of computer vision to compare surfaces [2–4, 7, 31, 41, 44]. Through spectral geometry, the geometry and topology of a shape is represented by its spectrum, which are the eigenvalues of the Laplace-Beltrami operator. The HKS, WKS and PLO3S descriptors are constructed with the eigenvalues and eigenfunctions of the Laplace-Beltrami operator.

These spectra have multiple properties, the central one being the invariance to isometry [2, 31, 44]. Invariance to isometry prevents high-amplitude, non-rigid transformations to modify the values of the descriptor, which is an important feature to take into account for protein shape comparison since proteins are dynamic objects that display different conformations.

To our knowledge, no method has been designed to find local surficial similarities of proteins independently of the sequence using the spectra of the Laplace-Beltrami operator. The Surface Wave Interpolated Maps (SWIM) descriptor has been developed on a protein surfaces dataset and is based on the spectra of the Laplace-Beltrami operator through the Wave Kernel Signature (WKS) descriptor. The SWIM descriptor also reduces the dimensions to a 2D space, which reduces the computation time compared to classic spectra-based methods used for 3D objects comparison such as HKS and WKS.

Although SWIM is a 2D descriptor and some information is lost when the dimensions are reduced, the precision-recall curves indicate a better performance of PLO3S compared to the other computer vision methods. Protein surfaces are rough, displaying many variations over their surface. Since computer vision methods are designed to be applied on objects that often display flat surfaces, the rough protein surfaces is considered as a noisy signal which decreases their performance in retrieval. In PLO3S, this problem is overcome by smoothing the surface shape (an average of the points of the 3D surface is projected on the same point on the 2D unit sphere).

For a set of *n* objects, an all-against-all comparison requires *n × n* comparisons, while only *n* descriptor computations are required (once per object). For this reason, the speed of the descriptors comparison affects the method time to a higher degree than the descriptor computation time. The slow computation of SWIM compared to the computation of HKS and WKS descriptors is not an issue in this context since it is a one-time operation that can be pre-processed. On the contrary, the computation of dissimilarity in PLO3S is 272 times faster than in WKS and 344 times faster than in HKS. This is due to (1) the definition of SWIM in the 2D space (2) the creation of a specific data structure that can be manipulated by the GPU with GPGPU. To the best of our knowledge, there is no GPGPU implementation of HKS and WKS.

The performances of PLO3S in enrichment are similar or higher than the reference computer vision methods evaluated in the present work with a faster comparison time to compute the dissimilarity. PLO3S can be used to quickly and efficiently decrease the size of a very large dataset while retaining the real positive objects. This method can be used for a screening in a big data environment to reduce a protein dataset that can be refined with a finer-grained method in a hierarchical protocol.

PLO3S is a local protein surface comparison method independent based on a surface descriptor that is solely based on the surface shape, then independent of the sequence, structure or fold of the protein.

Since proteins with a related function often share a similar surface while potentially displaying a low sequence, structure or fold similarity [15, 23, 39], PLO3S can be used in complement to protein structure comparison methods in different applications where the identification of protein surficial homologs is important such as target fishing for adverse interaction screening or poly-pharmacology in a drug discovery pipeline and protein-protein interactions annotation.

## 5 Conclusion

In the present work, we introduce PLO3S that relies on a 2D representation of the surface topology based on a conformal projection of the protein surface, the SWIM descriptor. The SWIM descriptor allies the advantages of being spectrum-based, *i.e* invariant to isometry and accounting for local surficial features of the shape, and of being in 2D, allowing very fast computation of the shape dissimilarity for the screening of large protein surface datasets. In addition, SWIM is a local descriptor that allows for a partial comparison and can therefore be used to find similarities between protein regions.

The performance of PLO3S in enrichment has been evaluated in a blind comparison of protein surfaces using the SHREC 2020 protein shapes database. The PLO3S method can be used as a fast, coarse grained protein surface shape screening method that efficiently eliminates proteins displaying dissimilar surfaces to downsize large datasets of protein surface shapes.

Since proteins with a related function often share a similar surface while potentially displaying a low sequence, structure or fold similarity [15, 23, 39], PLO3S can be used in complement to protein structure comparison methods in different applications where the identification of protein surficial homologs is important such as target fishing for adverse interaction screening or poly-pharmacology in a drug discovery pipeline and protein-protein interactions annotation.

## Acknowledgment

This work has received funding from the European Research Council (ERC) under the European Union’s Horizon 2020 research and innovation programme (grant agreement n° 640283).

